# Non-Parametric Rank Statistics for Spectral Power and Coherence

**DOI:** 10.1101/818906

**Authors:** Bahman Nasseroleslami, Stefan Dukic, Teresa Buxo, Amina Coffey, Roisin McMackin, Muthuraman Muthuraman, Orla Hardiman, Madeleine M. Lowery, Edmund C. Lalor

**Affiliations:** Academic Unit of Neurology, Trinity College Dublin, the University of Dublin, Dublin, Ireland; Biomedical Statistics and Multimodal Signal Processing Unit, Department of Neurology, University Medical Center of the Johannes Gutenberg-University, Mainz, Germany; Trinity College Institute of Neuroscience, Trinity College Dublin, the University of Dublin, Dublin, Ireland; Beaumont Hospital, Dublin, Ireland; School of Electrical and Electronic Engineering, University Colelge Dublin, Dublin, Ireland; Trinity Centre for Bioengineering, Trinity College Dublin, the University of Dublin, Dublin, Ireland; Departments of Biomedical engineering and Neuroscience, University of Rochester, Rochester, NY, USA

## Abstract

Despite advances in multivariate spectral analysis of neural signals, the statistical inference of measures such as spectral power and coherence in practical and real-life scenarios remains a challenge. The non-normal distribution of the neural signals and presence of artefactual components make it difficult to use the parametric methods for robust estimation of measures or to infer the presence of specific spectral components above the chance level. Furthermore, the bias of the coherence measures and their complex statistical distributions are impediments in robust statistical comparisons between 2 different levels of coherence. Non-parametric methods based on the median of auto-/cross-spectra have shown promise for robust estimation of spectral power and coherence estimates. However, the statistical inference based on these non-parametric estimates remain to be formulated and tested. In this report a set of methods based on non-parametric rank statistics for 1-sample and 2-sample testing of spectral power and coherence is provided. The proposed methods were demonstrated and tested using simulated neural signals in different conditions. The results show that non-parametric methods provide robustness against artefactual components. Moreover, they provide new possibilities for robust 1-sample and 2-sample testing of the complex coherency function, including both the magnitude and phase, where existing methods fall short of functionality. The utility of the methods were further demonstrated by examples on experimental neural data. The proposed approach provides a new framework for non-parametric spectral analysis of digital signals. These methods are especially suited to neuroscience and neural engineering applications, given the attractive properties such as minimal assumption on distributions, statistical robustness, and the diverse testing scenarios afforded.

## 1 Introduction

The frequency domain analysis of time series, including the analysis of spectral power and the coherence between two (or multiple) signals (Brillinger, 2001) is pivotal to the analysis of neural signals. The spectral analyses of neuro-electric signals such as electroencephalographic (EEG) and electromyographic (EMG) signals have been specifically of interest (Rosenberg, Halliday, Breeze, & Conway,1998; D. Halliday et al., 1995; D. M. Halliday & Rosenberg, 1999), as they reflect the intensity of neural oscillations in the nervous system, as well as the short and long range communications in the neural circuits through the oscillation synchrony or coherence (Siegel, Donner, & Engel, 2012). The discovery and quantification of fundamental neurophysiological phenomena such as cortico-cortical coherence (Walter, 1963; Koles & Flor-Henry, 1987), intermuscular coherence (Person & Kudina, 1968; Farmer, Bremner, Halliday, Rosenberg, & Stephens, 1993) and cortico-muscular coherence (Conway et al., 1995; D. Halliday et al., 1995) and subsequent contributions to the study of motor system neurophysiology (Boonstra, 2013; Wijk, M, Beek, & Daffertshofer, 2012) and pathophysiology (K. M. Fisher, Zaaimi, Williams, Baker, & Baker, 2012; Nasseroleslami et al., 2017) have been underpinned by spectral power and coherence analysis of the neuroelectric signals.

Despite advances in the application of spectral power and coherenceanalysisofneuralsignalsin (motor) neuroscience (Nasseroleslami, Lakany, & Conway, 2014; Waldert et al., 2009), neural and rehabilitation engineering (Aleksandra Vuckovic et al., 2014; Xu et al., 2014; A. Vuckovic et al., 2015) and neuro-diagnostics (Iyer et al., 2015; Nasseroleslami et al., 2017), the statistical inference of these measures in practical and real-life scenarios remains a challenge. The high dimensional nature of EEG and EMG signals that span a few to hundreds of (spatial) channels, different frequency-bands and different time windows is a challenge that can be addressed by using contracted/abstract measures (e.g. band-specific averages, number of detections and maximum/peak value and locations), or more systematically by false discovery rate corrections (Benjamini & Hochberg, 1995; Benjamini, Krieger, & Yekutieli, 2006) and empirical Bayesian inference (Efron, Tibshirani, Storey, & Tusher, 2001; Efron, 2004, 2007). There are, however, more fundamental challenges in statistical inference of spectral measures of neuro-electric signals. First, the statistical distribution of the neural signals can deviate from normal distribution, due to the complex underlying neurophysiological phenomena (Nasseroleslami et al., 2014; Mehrkanoon, Breakspear, & Boonstra, 2014), non-stationarity (A. Anwar et al., 2014; van Ede, Quinn, Woolrich, & Nobre, 2018), and other potentially unknown factors. Moreover, presence of various artefactual components (Tatum, Dworetzky, & Schomer, 2011) lead to extreme outliers (Nolan, Whelan, & Reilly, 2010) that lead to non-normal distributions and dramatical bias of the parametric measures of centrality. Artefact rejection methods (Mohr, Nasseroleslami, lyer, Hardiman, & Lalor, 2017; Nolan et al., 2010) can reduce some (but not all) of these unwanted effects due to artefacts, but not those originating from the underlying neurophysiological complexity and the inherent non-stationarity or non-linearities. These issues are major obstacles against using the existing parametric methods (Brillinger, 2001; Bloomfield, 2004; Priestley, 1982) for inferring the presence of specific spectral components above the chance level (1-sample testing) or presence of a significant differences between conditions or sets of signals (2-sample testing).

We have recently shown that using the median rather than mean for estimating the auto-spectra, cross-spectrum, and coherence can attenuate the effect of artefactual components in EEG connectivity analysis (Dukic et al., 2017). However, the analysis of the statistical significance of non-parametric spectra and coherence measures can no longer be assessed using the traditional estimates for the expected values of spectra that rely on the mean operator (Rosenberg et al., 1998; D. Halliday et al., 1995; D. M. Halliday & Rosenberg, 1999). Non-parametric rank statistics (e.g. Sign Tests, Wilcoxon’s Signe dRank Test, Mann-Whitney U Test, and Kruskal-Wallis Analysis of Variance) have provided promising approaches to statistical inference (Akritas, Lahiri, & Politis, 2014; Puri & Sen, 1971). However, these methods have not been translated to spectral signal analysis. In this report a new non-parametric approach for estimating the spectral power and coherence measures is explained and new methods based on non-parametric rank statistics are developed in order to assess the significance of the power and coherence, as well the differences between the levels of two power or coherence measures. We will demonstrate and test the utility of these measures through simulated signals and provide an example on the application of these measures in the context of research on neuro-electric signals.

## 2 Original Formulation for Spectra and Coherence

Consider *x*(*t*) and *y*(*t*) to be time domain signals (with *t* representing time), and *x_i_*(*t*), *y_i_*(*t*)representing one of the *L* epochs of the signals. The frequency domain representation of the signal, i.e. the (Discrete) Fourier Transform of each epoch is then represented by the complex-valued *X_i_*(*f*) and *Y_i_*(*f*) (*f* is the frequency). The auto-spectral densities *F_xx_* and *F_yy_*, as well as the cross-spectral density *F_xy_* are defined as (Bloomfield, 2004; Brillinger, 2001):

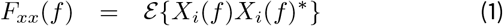

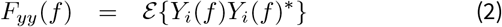

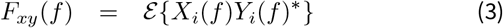

where * denotes the complex conjugate and 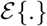 denotes the expected value, which is typically taken as the arithmetic mean of the values across the Lepochs, i.e.

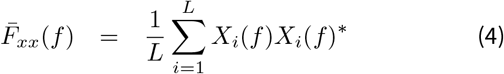

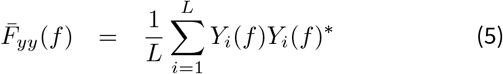

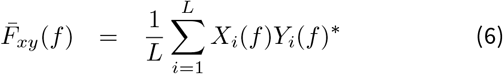

The bar symbol (^−^) here indicates that the expected value was found using the mean operator.

The standard deviation of auto-spectra can at frequency *f* be approximated using the variance *var*{*F_xx_*(*f*)} ≈ (*F_xx_*(*f*))^2^/*L*. In practice, the more useful relationship is that of the signal’s asymptotic variance. Therefore, *log_e_*(*F_xx_*(*f*)) will have the standard deviation of 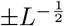 (Bloomfield, 2004; D. Halliday et al., 1995). This standard deviation (or variance) value can be used to identify statistically significant oscillations. Specific spectral components are significantly different from this signal value as tested by a 1-sample t-test (or a z-test). The log-transformed parametric variances are, however, may not be stable and suitable for 2-sample testing of the levels of spectral power in 2 different conditions.

The spectral coherency function is defined as a function of the auto-spectra and cross-spectrum, and therefore given by:

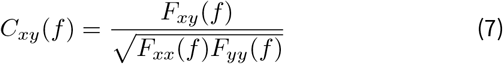

The coherence, defined as |*C_xy_*(*f*)|^2^ is a commonly used measure to assess the frequency domain correlation or phase synchrony between *x*(*t*) and *y*(*t*). It can be rewritten and interpreted as a measure of phase synchrony across epochs that is weighted by the magnitude values (Cohen, 2014). Statistical analysis of coherence is complicated in the general form, as it has a hyper-geometric distribution (Priestley, 1982). However, for the null hypothesis (where the coherence is 0), the distribution can be very accurately approximated. In this case a *tanh*^−1^(.) transforms the null coherence distribution to normal (Brillinger, 2001; Bloomfield, 2004; D. Halliday et al., 1995). Therefore for a given significance level *α*, the one-tail threshold value for significant coherence would correspond to 1 − *α*^1/(*L*−1)^, or:

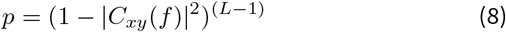

Equation (8) can be used for hypothesis testing against a significance threshold *α* such as 0.05 or 0.001. This estimate is very accurate under the normal distribution assumption of the time domain signals, and is a powerful statistical test. However, the 2-sample testing of 2 different coherence values, possibly with the ability to detect changes in both amplitude and phase is not adequately established.

## 3 Non-Parametric Estimation of Spectra and Coherence

We have recently demonstrated (Dukic et al., 2017) that in order to reduce the effects of artefactual components and account for non-normal distribution of signals, we may choose to use the Median operator as an estimator for the auto- and cross-spectra:

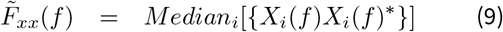

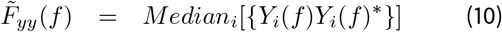

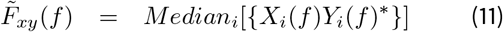

The tilde symbol (^~^) here indicates that the expected value was found using the median operator.

Figure 1 shows the statistical distribution of 2 exemplary raw auto-spectra, as well as the auto-spectra values estimated using the traditional mean operator, 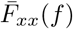 or the new median operator, 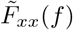. As the autospectra *F_xx_*(*f*) and *F_yy_*(*f*) are real-valued 1D variables, the definition on median, as the value of the inverse cumulative distribution function at 0.5, iCDF (0.5), is straightforward.

**Figure 1.**
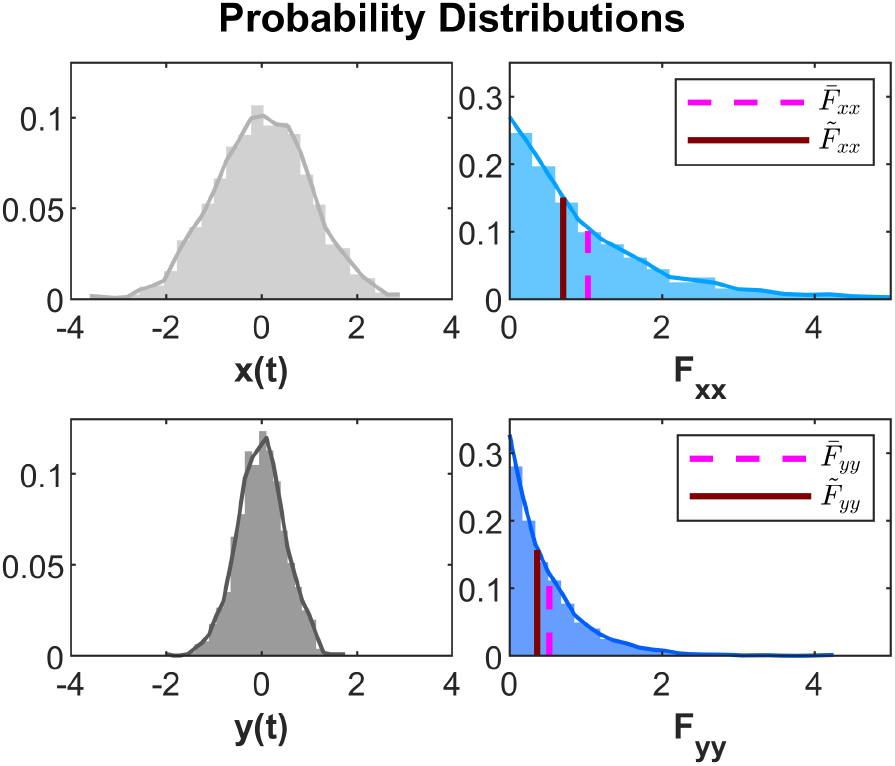
Statistical Distribution of the Time-Domain Signals, the Raw Auto-Spectra and The Estimation of Auto-Spectra using the Mean 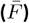 and Median 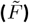. The simulated data are 2000 epochs of 1-s long random normal data (white noise) at 1000Hz for *x*(*t*). *y*(*t*) was taken as linear combination of similar random data with weight 0.2 and signal *x* with the weight 0.8.*e*^*jπ*/4^. The data used in histograms were taken from 10^th^ time points and frequency values at 10Hz..

Figure 2 shows the statistical distribution of an exemplary raw cross-spectra. The 2D distribution of the complex-valued raw cross-spectra, as well as the marginal distributions (projected to the real and imaginary axes, respectively) are shown. The 2D cross-spectrum values estimated using the traditional mean operator, 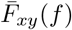 or the new median operator, 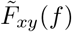 are shown on the 2D and marginal distributions.

**Figure 2.**
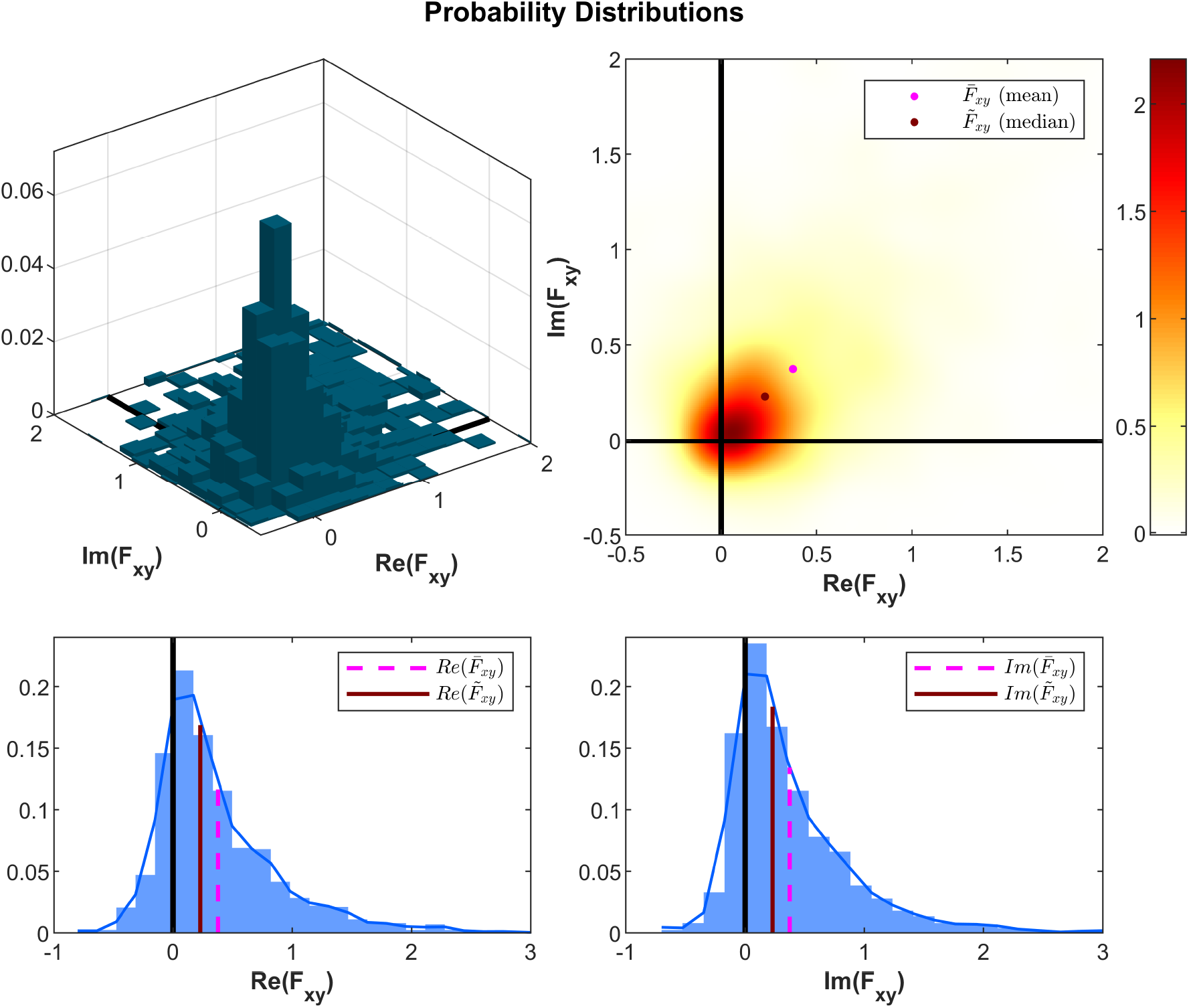
Statistical Distribution of the Raw Cross-Spectra in 2D and Marginal Plots and the Estimation of Cross-Spectrum using the Mean 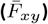 and Median 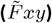. The 2D plots (top) show the probability distribution functions as a function of the real and imaginary components and the 1D plots (bottom) show the marginal distribution of the real and imaginary components in the presence of a non-zero coherence. The frequency values correspond to the 10Hz of the *x* and *y* signal as in Figure 1.

The definition of the 2D median can be based on several (slightly) different criteria (Pranab K. Sen, 2005; Niinimaa & Oja, 2006). The simplest estimate of 2D median is the marginal median, whose components are calculated as 1D median in each dimension of the multivariate data (Niinimaa & Oja, 2006). Therefore, the complex (marginal) median is found by combining the median of the real and the median of the imaginary values (Dukic et al., 2017):

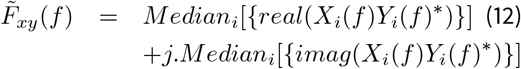

The other approach is to use the ***Spatial Median***, which is based on the minimum Euclidean distance from data points (Niinimaa & Oja, 2006).

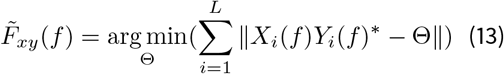

This is a more favourable choice, given its more useful properties in 1-sample and 2-sample tests.

Following the calculation of the non-parametric auto-spectra and cross-spectrum, the coherence may be estimated similar to the parametric case:

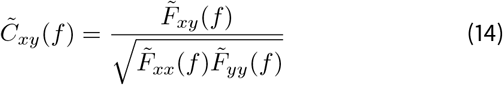

This non-parametric coherency, usually described in terms of the magnitude 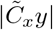, can be interpreted similar to the parametric definition 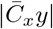. However, both measures are biased and the values that correspond to specific significance levels depend on the number of epochs *L* used for calculation, as well as other potential spectral smoothing procedure. Therefore, to account for these concerns and to elevate the numerical instability of 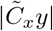 found by small number of epochs *L*, the spectral power and coherence measures are best described in terms of their p-values. This will be established in the following section.

## 4 Non-parametric Rank Statistics for Spectral Power

The statistical distribution of the raw auto-spectra {|*X_i_*(*f*)|^2^}can be used to define specific percentiles of the distributions using the inverse cumulative distribution functions, iCDF. For example, the 2-tail 95% confidence interval with *α* = 0.05 would be:

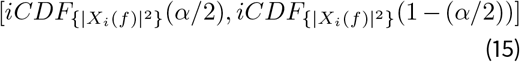

which is defined non-parametrically on the signal values at the desired frequency or band *f*.In case the sampling distribution (commonly needed for statistical testing) of the measures 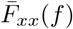 or 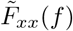 are required, they can be found by bootstrapping; however, this is not primarily of interest, as the statistics will be handled using non-parametric rank-based methods.

### 4.1 1-Sample Power: Testing for Significant Spectral Activity

A hypothesis that is more commonly needed to be tested is the presence of a neural oscillations (spectral power) at a specific frequency (band), beyond that of other frequencies in the null situation (e.g. a white noise). The above-mentioned confidence intervals may prove useful. However, in practical applications such as in neural signal analysis it is usually desired to establish the significant presence of activity in specific frequencies with respect to the others. The following procedure explains a method for such a statistical testing.

If |*X_i_*(*f_j_*)|^2^ shows the power pertaining to the epoch *i* (of total *L*), and the frequency value or frequency band *j* (of total *M* selected frequency bands for analysis, excluding 0 or DC values), we may find for each epoch *i*, the tie-adjusted rank *R_xx, i_*(*f_j_*) among the *M* values which will give values in the range [1, *M* − 1]. We can then use the centred version of the rank *R_xx, i_*(*f_j_*) – (*M*/2), and consequently apply a non-parametric test such as Wilcoxon’s Signed Rank test on the *L* centred rank values *R_xx, i_*(*f_j_*) – (*M*/2), *i* = 1…*L* which tests for higher or lower frequency content at the tested frequency *fj*. This procedure detects a significant presence (or absence) of spectral or oscillatory activity under the null assumption of the white noise distribution. The test can be modified for testing against different null hypotheses (e.g. a noise with 1/*f* distribution), by subtracting the profile of the null auto-spectrum from the {|*X_i_*(*f_j_*)|^2^} data before forming the rank values. Figure 3 provides a pictorial explanation of this testing procedure for detecting a sinusoidal component mixed with a white noise. It shows a median-based estimate of auto-spectrum 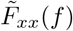, as well as the p-value of the spectrum associated with the oscillatory activities in the signal.

Notice that if the intention is to detect only the increase (or decrease) in spectral components a right-tail or left-tail test may be used.

**Figure 3.**
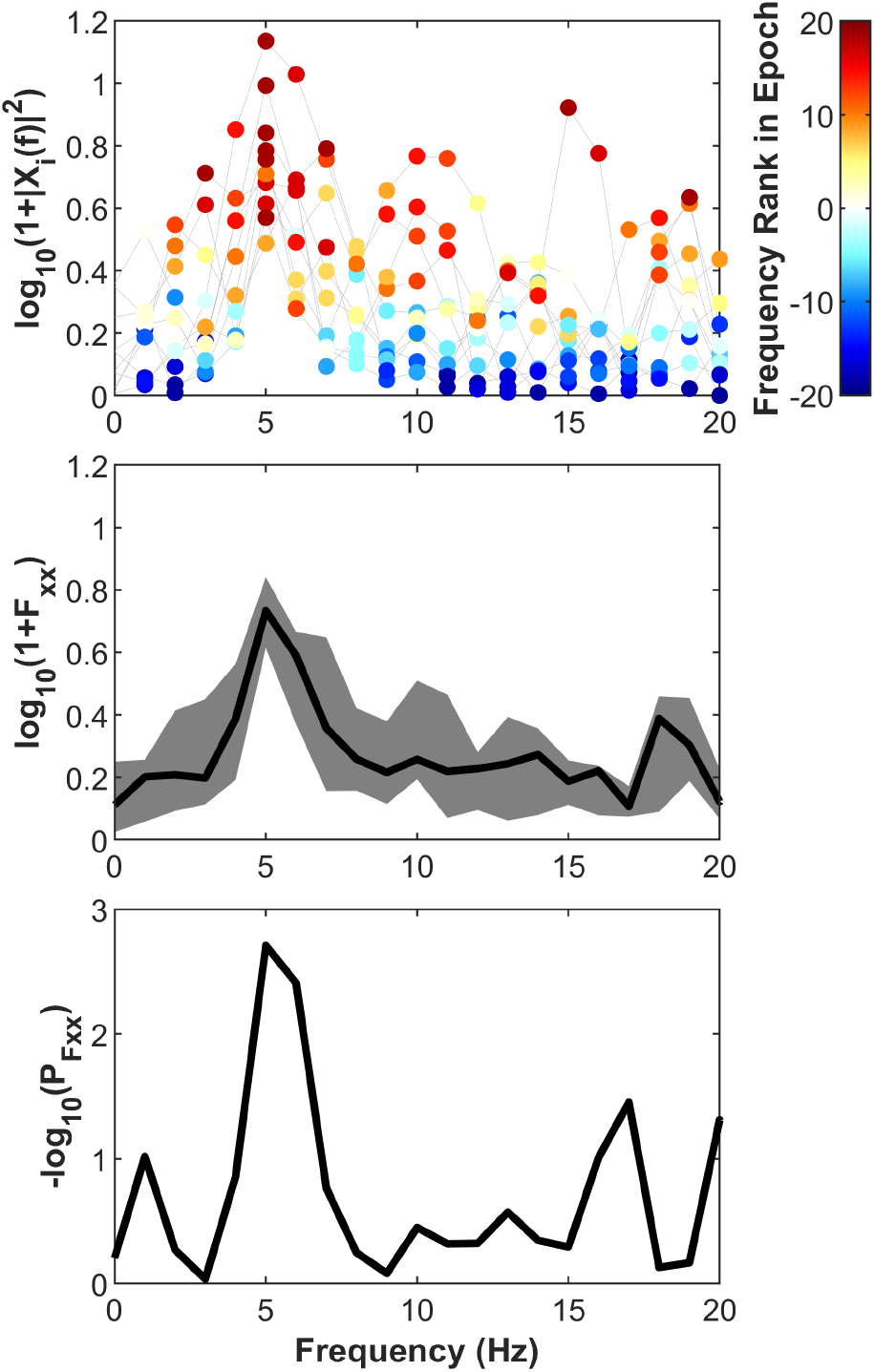
Finding the Significance of Oscillatory Power by Rank Statistics. **Top:** Rank ordering is first applied across frequencies at each epoch. The centred ranks are subsequently tested at each frequency using Wilcoxon’s Signed Rank test. **Middle:** Median-based estimate of autospectrum 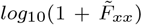 and the interquartile range of the raw auto-spectra **Bottom:** p-value of the spectral power against the null hypothesis of no significant presence of specific oscillatory components (whitenoise). Thedataforsimulationwas taken as 10 epochs of 40 sampled at 40Hz, a linear combination of white noise with the weight of 1 and a sinusoidal waveform at 5Hz with the weight of 0.85..

### 4.2 Sample Power: Testing for Significant Change of Power of Between Groups

Testing for between-group comparisons is straightforward as the data {|*X_i_*(*f*)|^2^}and {|*Y_i_*(*f*)|^2^} can be directly tested with Mann-Whitney U test. As the non-parametric two-sample location test can be applied on 1D data with non-normal distributions, the power values in individual epochs can be directly compared. Notice that the number of epochs for this comparison can be dissimilar.

## 5 Non-parametric Rank Statistics for Coherence

The cross-spectrum 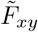 or the coherence 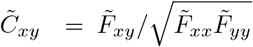 defined based on the medians of the auto- and cross-spectra can be tested against the null hypothesis. The test for presence of significant coherence is a 1-sample 2D location problem where the cross-spectrum is compared against 0 + *j*0. The test for comparing the coherence between 2 pairs of possibly coherent signals is a 2-sample 2D location problem. Each of these scenarios will be discussed as follows.

### 5.1 1-sample Coherence: Testing for Significant Coherence

The L complex values {*X_i_*(*f*)*Y_i_*(*f*)*}in the 2-Dimensional complex plane need to be compared to the point (0, 0). This is a 2-dimensional 1-sample location problem (Figure 4). A parametri version of such a test is the 1-sample Hotelling’s *T*^2^test (Hotelling, 1931). However, in addition to the parametric nature of the test, the 2D distribution of cross-spectrum data is not normal and is not straightforward to convert to normal by either functional (e.g. log(.)) (Box & Cox, 1964) or inverse normal transformations (Beasley, Erickson, & Allison, 2009). Therefore, in order to follow the non-parametric route, {*X_i_*(*f*)*Y_i_*(*f*)*} can be tested against zero using non-parametric 2-dimensional sign or rank statistics. The statistical tests based on marginal medians (Pranab Kumar Sen & Puri, 1967; Puri & Sen, 1971) are simple to use. However, they are not affine-invariant; therefore, in the context of spectral analyses they would produce variable results for the same level of coherence but with different phase differences (corresponding to a rotation transformation in the complex plane). Therefore, only affine-invariant tests are suitable for this 2D location problem. An example of these tests are the tests based on spatial median and ***spatial (signed) ranks*** (Hannu Oja & Randles, 2004; Hannu Oja, 2010; Nordhausen & Oja, 2011). In R (R Core Team, 2016), the R-package MNM (Nordhausen & Oja, 2011; Nordhausen, Mottonen, & Oja, 2018), this test has been implemented as “*mv.1sample.test*”, a signed-rank based test with inner standardization. Notice that this 1-sample test in fact corresponds to the presence of significant magnitude. A separate non-parametric test to compare the phase of the coherence against a hypothesised value *ϕ*_0_ can be achieved by applying the Wilcoxon’s Signed Rank test on the data {∠(*X_i_*(*f*)*Y_i_*(*f*)*) – *ϕ*_0_} (represented in the range [−*π, π*]).

**Figure 4.**
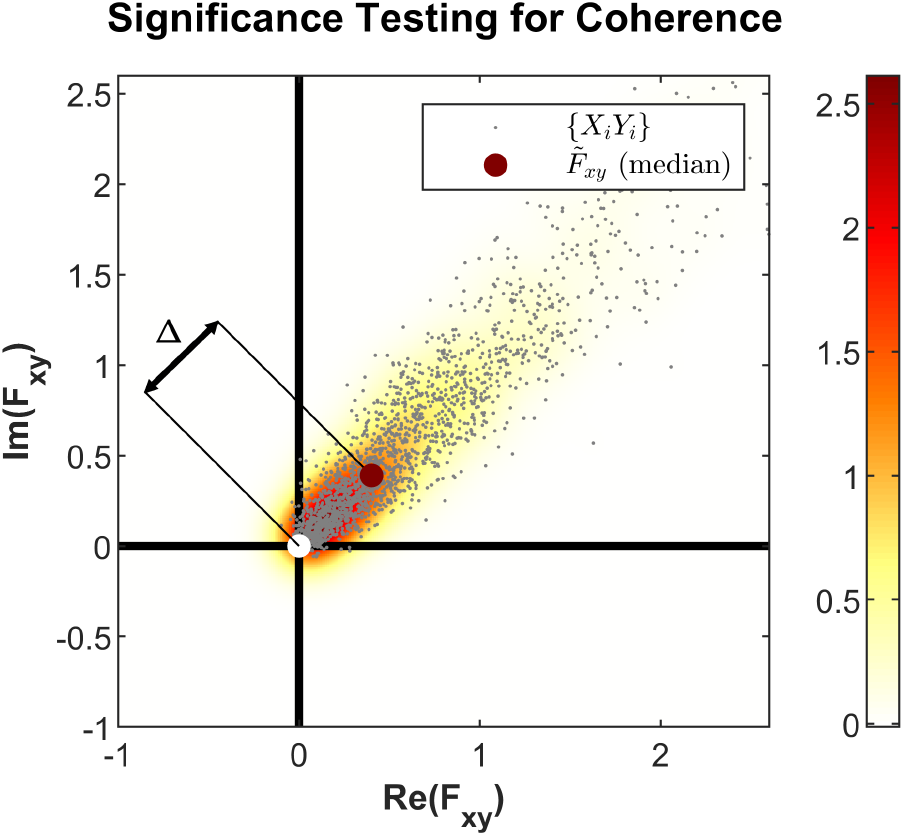
Non-Parametric 1-Sample Testing for Significant Coherence.. The dots show the complex-values raw cross-spectra data points and the colour shows the 2D probability density function for the same data. The raw cross-spectra (i.e. the median-based estimate of coherence 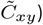 are compared against the zero, i.e. the origin (0, 0). This corresponds to testing for the difference between the coherence magnitude 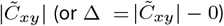 against zero. Notice that as the the division by 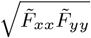 is only a scaler scaling factor in the definition of coherence in Equation (14), the comparison against zero for cross-spectrum and coherence are equivalent. The simulated data are similar to the description in Figure 1, taken at 10Hz.

### 5.2 2-Sample Coherence: Testing for Significant Difference of Coherence Between Groups

In 2-sample problems, it may be desired to compare the coherence between the 2 signals *x*(*t*) and *y*(*t*) in an experimental condition 1 with *L*_1_ epochs against their coherence in experimental condition 2 with *L*_2_ (or to compare the synchrony of 2 different pairs of signals). The existing methods for comparing 2 levels of coherence values in 2 groups has limitations. One limitation is the inability to simultaneously test for a difference in magnitude and phase of coherence, with no solid method to test for phase difference. Importantly, comparisons at individual subject levels are not established.

#### 5.2.1 Comparison of Phase

The comparison of phase between the spectral data {*X_i_*(*f*)*Y_i_*(*f*)*|*i* = 1…*L*_1_} (with coherence 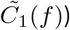 and {*X_k_*(*f*)*Y_k_*(*f*)*|*k* = 1…*L*_2_} (with coherence 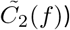 can be achieved using the circular statistical tests. The parametric and non-parametric circular tests for comparing {∠(*X_i_*(*f*)*Y_i_*(*f*)*)}and {∠(*X_k_*(*f*)*Y_k_*(*f*)*)}would be the Watson-Williams (Watson & Williams, 1956) and Fisher’s (N. I. Fisher, 1995) tests. Such tests provide valid results only under certain assumption (e.g. on the presence of a minimum circular clustering in data), therefore they may not provide reliable estimates and a uniform p-value distribution under the null condition. An alternative simple approach is to find the circular mean of the data in both samples and then find the (minimum) circular distance of the data in each group to this common circular mean. The distances are consequently compared using the 2 sample t-test (as the parametric case) with unequal variances assumption. Given that the circular data are not actually normally distributed (nor necessarily have a Von-Mises-like circular distribution), the non-parametric option to compare the phase values would be to apply the Mann-Whitney U test on the previously found distances in each group.

However, importantly, the results of such tests for 1D phase comparisons are limited, as the they are not weighted by the magnitude of the raw cross-spectra and their validity is conceptually questioned when one or both of the coherence values are very low and non-significant.

#### 5.2.2 Comparison of Magnitude

The comparison of the magnitude of coherence between the spectral data {*X_i_*(*f*)*Y_i_*(*f*)*} and {*X_k_*(*f*)*Y_k_*(*f*)*} is not straight-forward, due to the effects of phase in shaping the resultant coherence magnitudes. Consequently, only the resultant coherence values 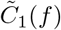 and 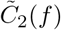 are available for testing, but not the accompanying standard deviations or individual data points. Nevertheless, under the assumption of equality of the phases between two coherent conditions (or its lack of importance), a statistical trick may be used for a quick test of difference between the strength of coherences. First both coherence values are compared against the null condition, i.e. two 1-sample tests are performed separately in each group and the resulting p-values *p*_1_ and *p*_2_ are found by either a parametric or non-parmetric test as described by Equation (8) or in Section 5.1. The p-values are then transformed to z-scores *z*_1_ and *z*_2_ by 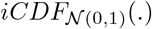, subtracted from each other and then difference Z-score is significance tested according to the resultant variance of the 2 z distributions, i.e 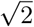. In other words:

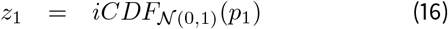

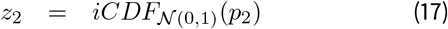

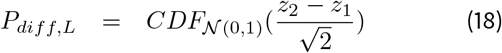

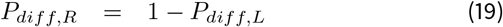

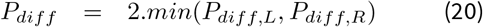

where 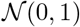 denotes the normal distribution with mean 0 and standard deviation 1; (i) CDF stands for (inverse) cumulative distribution function. The *P_diff, L_* and *P_diff, R_* are the left- and right-tail difference p-values and *P_diff_* the two tail p-value of difference between the 2 coherence levels. This procedure is similar to the Stouffer’s method (Stouffer, 1977; Westfall, 2014) for combining 2 or more p-values, except that the two z-scores are subtracted (rather than summed).

#### 5.2.3 Simultaneous Comparison of Magnitude and Phase

Using an approach similar to the 1-sample problem (See section 5.1), we may use 2-dimensional 2-sample location problems by considering the spectral data {*X_i_*(*f*)*Y_i_*(*f*)*} and {*X_k_*(*f*)*Y_k_*(*f*)*} as data points in the 2-dimensional real-complex plane (Figure 5). The difference can be tested parametrically using the 2-sample Hotelling’s *T*^2^test. This test, however, may not be preferred, given the sensitivity of test to the numerically calculated covariance matrices, which will be accentuated by the non-normal distribution of data and presence of artefacts. Similar to the 1-sample case, the 2-dimensional non-parametric tests can be used for this comparison. A simple and computationally efficient test is the Jurecková-Kalina (JK) test (Jurečková & Kalina, 2012) or two-sample tests for marginal medians (Puri & Sen, 1971). However, in order to achieve affine invariance property (as for the 1-sample case), tests based on spatial median and ***spatial ranks*** (Hannu Oja & Randles, 2004; Hannu Oja, 2010; Nordhausen & Oja, 2011) are preferred. The MMN R-package (Nordhausen & Oja, 2011; Nordhausen et al., 2018), has implemented this test as “*mv.Csample.test*”, a rank based test with inner standardization. Notice that this 2-/multi-sample test can in fact quantify the location difference between the 2 coherence levels, which may originate from differences in the magnitude, phase or both.

**Figure 5.**
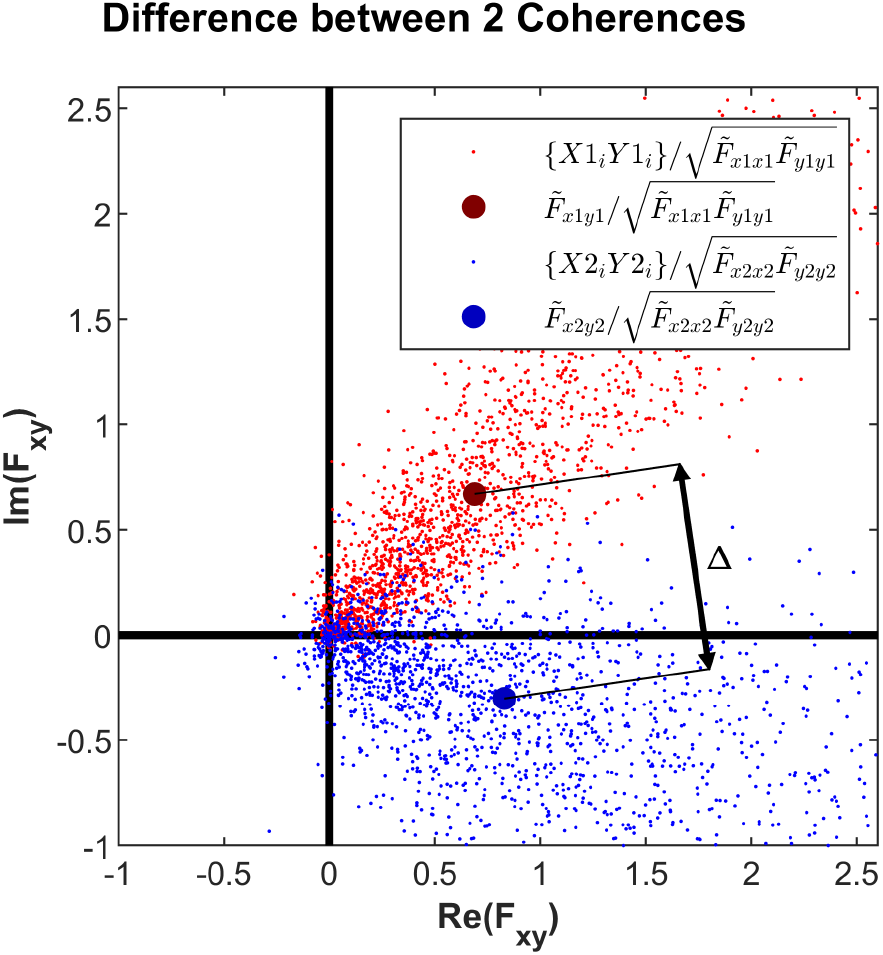
Non-Parametric 2-Sample Testing for Significant Difference between 2 Coherence Values. The dots show the complex-valued raw normalised cross-spectra data points for 2 pairs of signals (*x*_1_, *y*_1_) (red) and (*x*_2_, *y*_2_) (blue). The difference of distant between the raw normalised cross-spectra (i.e. the median-based estimates of coherence values 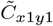 and 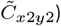 is compared against the zero. This corresponds to testing for the magnitude of difference between the coherence values 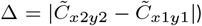 against zero. The simulated data are similar to the description in Figure 1, taken at 10Hz. *x*_1_ and *x*_2_ are constructed as white noise signals including 2000 epochs of 1s at 1000Hz. *y*_1_ in built by a linear combination of white noise noise and *x*_1_ with a factor 0.8.*e*^*jπ*/4^. *y*_2_ in built by a linear combination of white noise noise and *x*_2_ with a factor 0.7.*e*^−*jπ*/8^.

## 6 Numerical Examples

Several demonstrative examples are provided using both simulated data, as well as real-life examples using experimental EEG and EMG data.

### 6.1 Simulated Data

Before generating the demonstrating examples below, the sanity of all of the tests were verified by ensuring that under the null condition the probability density function of the p-values are uniform.

In all the simulations, epochs of normally distributed random numbers (white noise) were considered as *x*(*t*) and *y*(*t*) signals. The presence of coherence was simulated by adding a sinusoidal waveform to the data in all the epochs, which constituted a ratio *r* of the combined total signal. A second condition for non-normal distributions was considered, where contamination by artefactual components yield nonnormal distributions (referred to here as the condition with artefacts). This latter condition was emulated by adding large white noises with non-zero baselines to 10% of the epochs. The p-values, expressed as −*log_10_*(*p*) were compared across the normal signal condition and the non-normal contaminated signal, when tested by both the commonly-used parametric tests, as well as the newly proposed use of non-parametric tests. Each condition was simulated several times to afford stable estimates of the average values for −*log*_10_(*p*).

#### 6.1.1 1-Sample Spectral Power

Figure 6 (top) shows the simulation results of a sinusoidal waveform (signal), mixed with a background white noise (noise). As the Signal-Ratio *r*, defined by the percentage of the amplitudes, increases from 0 to higher values, the detection of significant oscillatory component, quantified by the p-values of the parametric and non-parametric tests are shown. The −*log*_10_(*p*) increases as function of *r* for both the parametric and non-parametric tests. While the parametric test is more sensitive at higher *r* values, it is dramatically affected by artefacts. The non-parametric test showed a reasonable level of robustness against artefacts.

**Figure 6.**
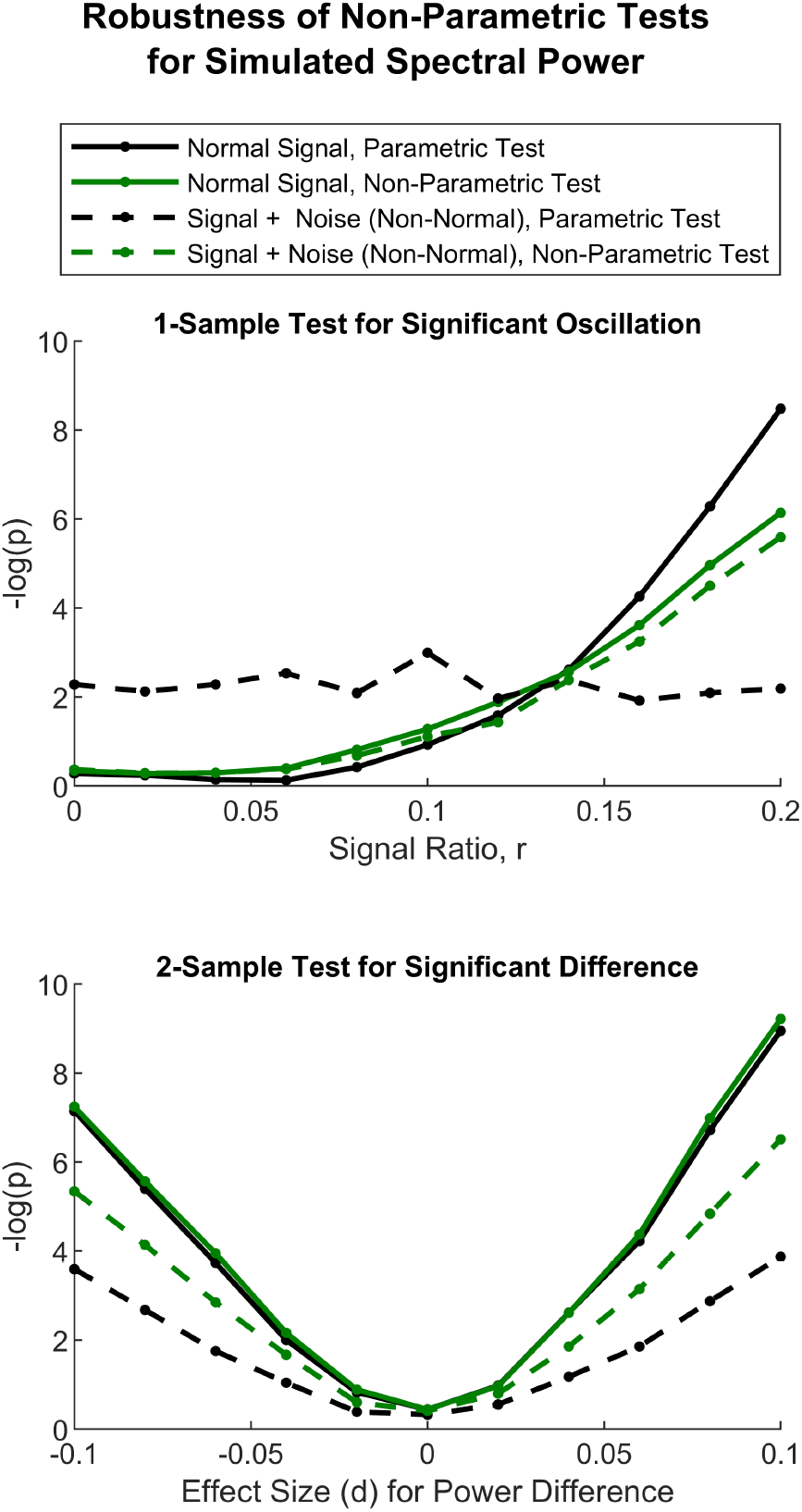
Simulation: Non-Parametric Statistical Tests of Spectral Power are Robust Against Artefacts in Simulations of 1-Sample and 2-Sample Comparisons. The simulated data include 50 epochs of random standard normal data (white noise) with 1s duration at 40Hz, combined linearly with a sinusoidal waveform at 10Hz with the relative signal weight *r*. For 1-sample test (top) the presence of the oscillatory component was detected in the signal combined with noise at different values of signal ratio *r*. For 2-sample tests (bottom), different levels of effect size *d* were tested by choosing *r*_1_ = 0.3 for *x*_1_ and varying *r*_2_ values for *x*_2_, so that *d* = *r*_2_ − *r*_1_ = *r*_2_ − 0.3. Artefactual components were added to 10% of epochs (randomly chosen) which were a combination of random white noise with standard deviations of 10, and a random shift between −5 and +5. All the tests were repeated 100 times and the average values are shown. The tests were performed on the values that correspond to 10Hz. See section 4.1 and Figure 3 for the details of 1-Sample non-parametric tests. For the 2-sample non-parametric tests, Mann-Whitney’s U test was used. For parametric tests, the z-scores, 1-sample and 2-sample t-tests replaced the centred ranks, Wilcoxon’s Signed Rank, and Mann-Whitney U tests, respectively.

#### 6.1.2 2-Sample Spectral Power

Figure 6 (bottom) shows the simulation results of 2 sinusoidal waveform (signals), mixed with background white noise (noise). As the effect size *d*, defined by the difference of the two signal ratios *r*_1_ = 0.3 and *r*_2_, increases from 0 to higher values, the detection of significant differences in spectral power are shown. The −*log*_10_(*p*) increases as function of *d* for both the parametric and non-parametric tests. The parametric and non-parametric tests result in very similar p-values as a function of the decrease and increase in the effect size. However, the non-parametric tests afford a considerably higher level of robustness against artefacts.

#### 6.1.3 1-Sample Coherence

Figure 7 shows the simulation results of 2 signals with *r*-percent shared synchronous sinusoidal oscillations. As *r* increases from 0 to 0.5, the detection of significant coherence is shown. The −*log*_10_(*p*) increases as function of *r* for both the parametric and non-parametric tests. While the non-parametric test is more sensitive at higher *r* values, it is dramatically affected by artefacts. The non-parametric test showed a reasonable level of robustness against artefacts.

**Figure 7.**
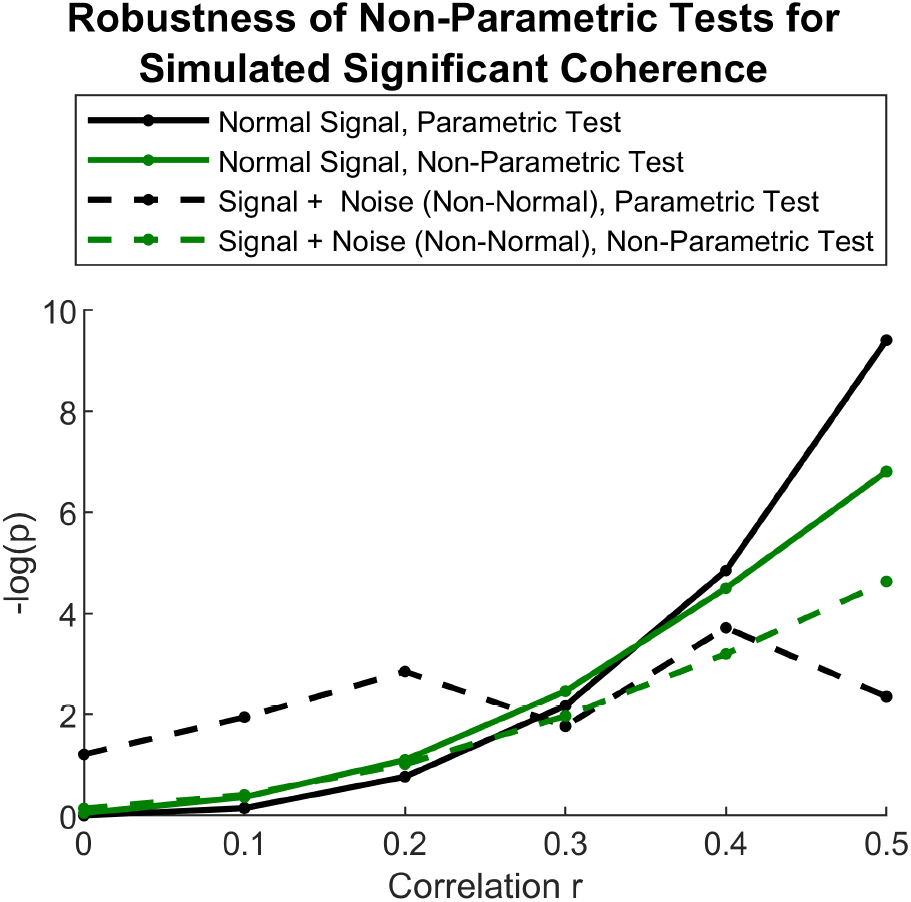
Simulation: The Non-Parametric 1-Sample Statistical Test, Spatial Signed Rank, for Presence of Significant Coherence is Robust Against Artefacts. The simulated data for *x*_1_ include 50 epochs of random standard normal data (white noise) with 1s duration at 40Hz. *y*_1_in built by a linear combination of white noise and *x*_1_ with a factor *r.e^jϕ^*. Artefactual components were added to 10% of epochs (randomly chosen) which were a combination of random white noise with standard deviations of 2, and a random shift between −2 and +2. All the tests were repeated 10 times and the average values are shown. The tests were performed on the values that correspond to 10Hz. The results shown correspond to phase difference *ϕ* = 0; the simulations with other phase difference values yielded the same results. See section 5.1 for details on the on-parametric Spatial Signed Rank Test and equation (8) for the description of the parametric test.

#### 6.1.4 2-Sample Coherence

To inspect the efficient detection of differences in 2 coherence levels, including both the magnitude and phase, 2 series of simulations were carried out using 2D tests. Each set of simulations assessed the effect of differences in either magnitude or the phase of coherence. In addition to comparisons between the parametric and non-parametric tests on normal and contaminated data, the 1D tests for comparing the magnitude or phase are provided as a reference.

##### Magnitude Difference

Figure 8 shows the simulation results of 2 pairs of signals with correlation values of *r* and 0.5 and the same phase values of coherence. As the difference between the correlation values changes from −0.3 to 0.3, the detection of significant difference between coherence levels is shown. The −*log*_10_(*p*) increases as function of difference for both the parametric and non-parametric tests. While the 2D parametric test is more sensitive than the 2D non-parametric test at higher *r* values, it is dramatically affected by artefacts. The 2D non-parametric test showed a reasonable level of robustness against artefacts. Similarly, the 1D parametric test is more sensitive than the 1D non-parametric test (and even the 2D parmetric test) at higher *r* values. It is, however, considerably affected by artefacts. The 1D non-parametric test showed a reasonable level of robustness against artefacts, but does not show a major advantage over the 2D non-parametric test except at lower coherence/difference values.

**Figure 8.**
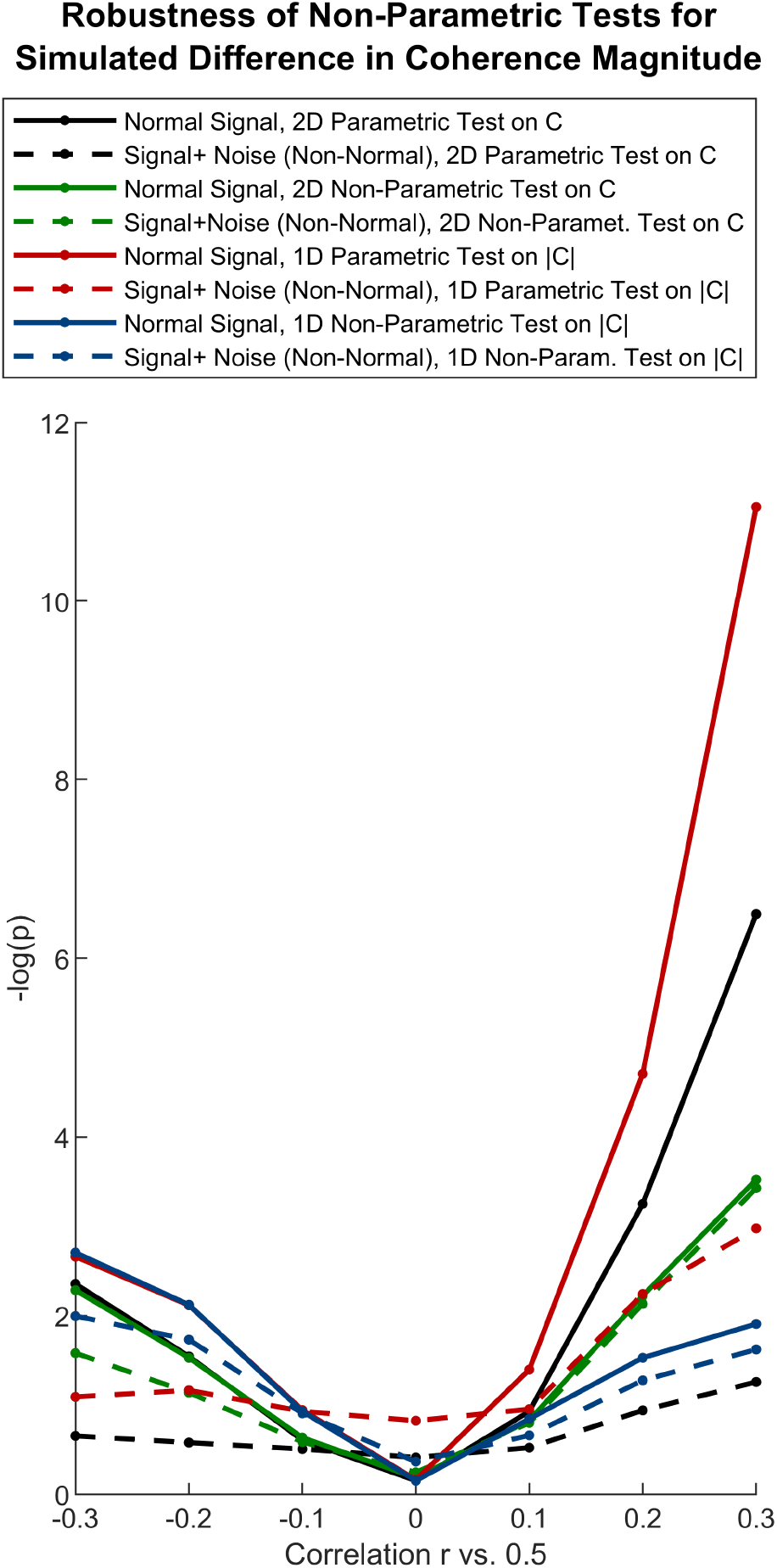
Simulation: The Non-Parametric 2-Sample Statistical Tests based on Spatial Rank and Spatial Signed Rank for Comparison of the Coherence Magnitude between 2 Conditions are Robust Against Artefacts. The 1D tests of difference use the statistical trick in section 5.2.2 for finding the difference based on 2 p-values (calculated parametrically from equation (8) or non-parametrically from 1-sample Spatial Signed Rank test in section 5.1). The 2D tests are 2-sample Hotelling’s *T*^2^ (parametric) and the 2-sample Spatial Rank test (non-parametric). Notice that the non-parametric tests are always more robust than parametric tests, where the 2D 2-sample test performs better at higher coherence values/differences and the 1D statistical trick on 1-sample non-parametric test (Section 5.1 and Figure 7) performs better on lower coherence values/differences. The simulated data for *x*_1_ include 50 epochs of random standard normal data (white noise) with 1s duration at 500Hz with weight 0.5, combined linearly with a sine waveform at 30Hz with weight 0.05. *y*_1_ is built similarly by a linear combination of white noise and sinusoidal waveform. *x*_2_ and *y*_2_ are similarly constructed, but with the weightings of 1 − *r* and 0.1*r* between the white noise and sinusoidal components. Artefactual components were added to 10% of epochs (randomly chosen) which were a combination of random white noise with standard deviations of 2, and a random shift between −2 and +2. All the tests were repeated 100 times and the average values are shown. The tests were performed on the values that correspond to 30Hz. The results shown correspond to phase difference *ϕ* = 0; the simulations with other phase difference values yielded the same results.

##### Phase Difference

Figure 9 shows the simulation results of 2 pairs of signals with similar correlation values of 0.5, but with coherence phase values changing systematically between [0, *π*]. As the difference between the phase values changes from 0 to *π*, the detection of significant difference between coherence values is shown. The −*log*_10_(*p*) increases as function of difference for both the parametric and non-parametric tests. While the 2D parametric test is more sensitive than the 2D non-parametric test at higher *r* values, it is dramatically affected by artefacts. The 2D non-parametric test showed a reasonable level of robustness against artefacts. Both the parametric and non-parametric 1-D tests for phase differences, performed poorly against the 2D tests.

**Figure 9.**
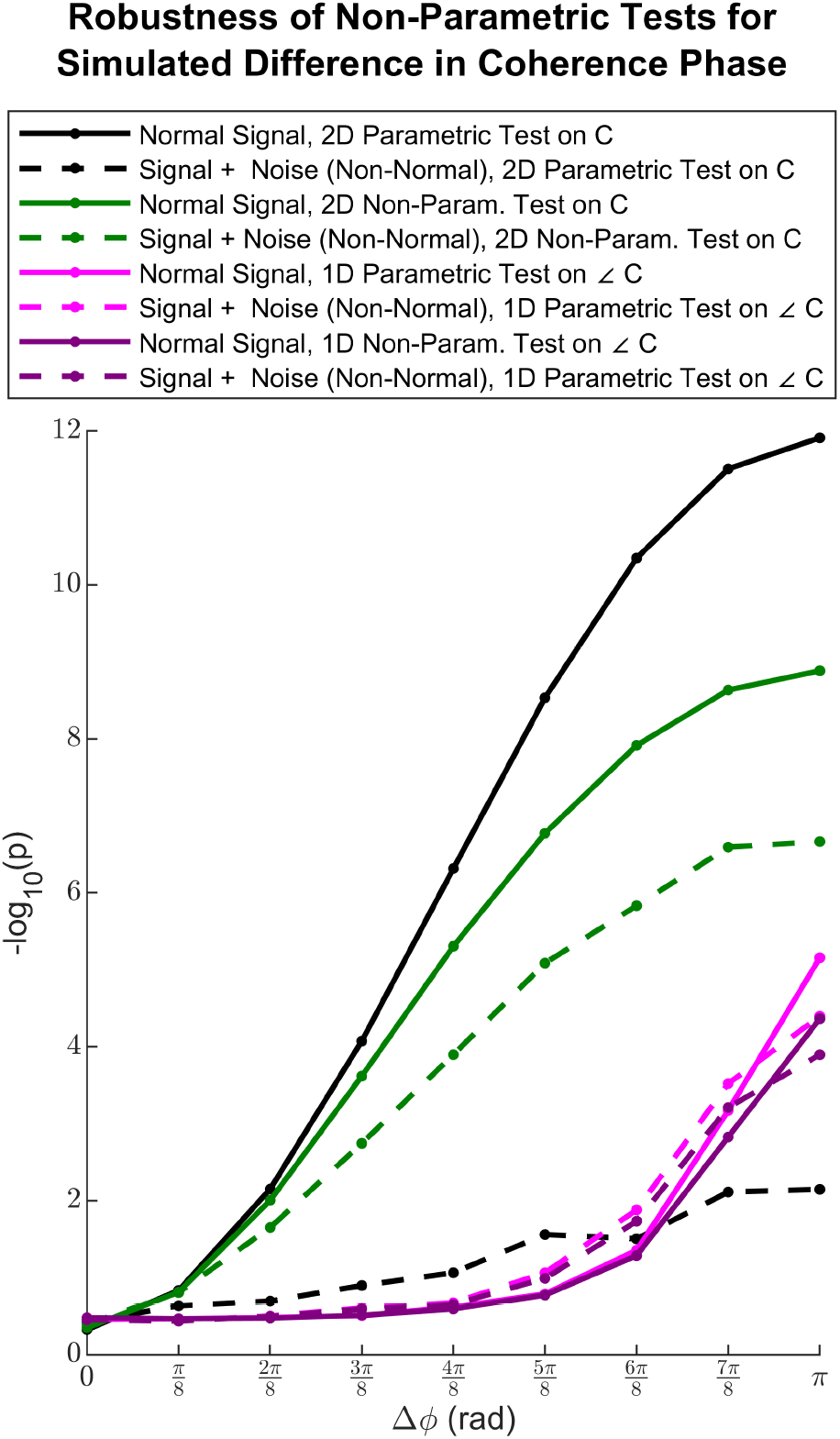
Simulation: The Non-Parametric 2-Sample Statistical Tests based on Spatial Rank and Spatial Signed Rank for Comparison of the Coherence Phase between 2 Conditions are Robust Against Artefacts. The 1D tests of difference use the circular 2 sample t-test (parametric) and circular Mann-Whitney U test (non-parametric) as described in section 5.2.1). The 2D tests are 2-sample Hotelling’s *T*^2^ (parametric) and the 2-sample Spatial Rank test (non-parametric). Notice that the 2D non-parametric test performs better than 1D tests and 2D non-parametric test and is robust. The simulated data is the same as data in Figure 8, with *r*_1_ = *r*_2_ = 0.5, the 0 phase between the sinusoidal components of *x*_1_ and *x*_2_ and phase difference of Δ*ϕ* between *x*_2_ and *y*_2_.

### 6.2 Experimental Data

The statistical tests described above were applied on real-life experimental data to further demonstrate their utility in practice. The experimental task was a sustained isometric pincher grip at 10% Maximum Voluntary Contraction (MVC) level. This was performed in 30 trials each lasting 5s. Experimental EEG and EMG data, including the Ear-Lobe-Referenced (ELR) EEG at electrodes *C*_3_ and *C*_4_, as well as bipolar EMG recorded from First Dorsal Interosseous (FDI) muscle were collected during the motor task. EEG and EMG signals were recorded at 2048Hz, filtered between 0 (DC) and 410Hz and stored for analysis. Data from a healthy control subject (age =?, gender=?) was used for analysis. A total of 102 epochs each lasting 1s that were correctly recorded were checked for artefacts (Dukic et al., 2017) and 99 acceptable epochs were used for analysis. Given the rejection of extreme artefacts, it is expected that both parametric and non-parametric provide comparable results.

#### 6.2.1 Spectral Power

Figure 10 shows the distribution of the raw autospectra for an EEG signal at different confidence intervals, the autospectrum calculated using the mean operator, as well as the auto-spectrum calculated using the median operator. Notice that even after the log-transform the spectrum has a non-normal distribution, as evidenced by the difference between the mean and median, as well as the poor prediction of the 68% CI by ±*σ*range. The p-values have similar overal behaviour, given that EEG signal is being tested against white noise as the null hypothesis. The parametric test is more effective and powerful at lower dominant frequencies, whereas the −*log*_10_(*p*) for non-parametric test is limited by the finite sample size and is saturated. On the other hand, non-parametric test performs better at frequencies about 400Hz, where the physiological content of the biosignals degrade, recording equipment filters part of the signal components, and the signal to noise ratio is very low.

**Figure 10.**
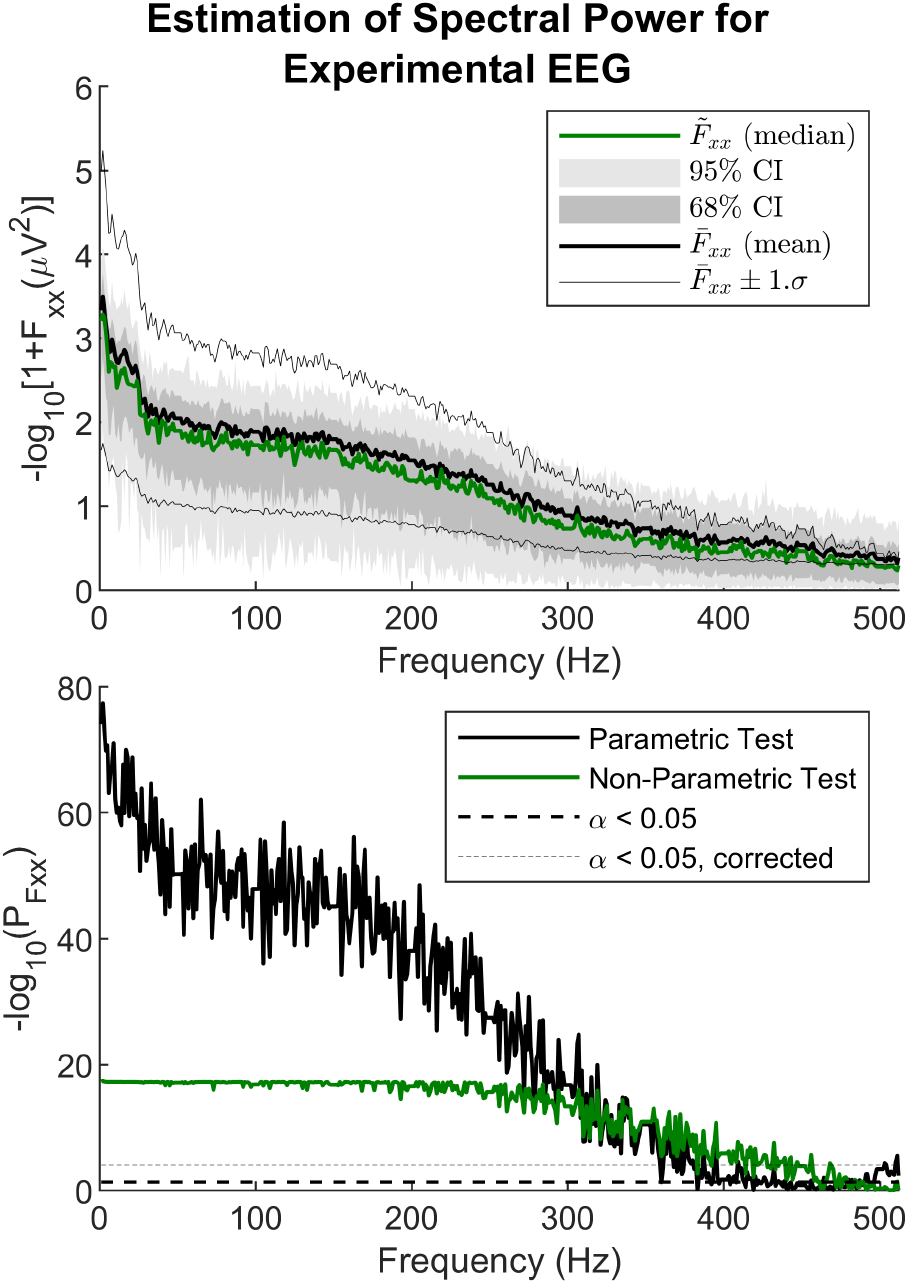
Experimental Data: Both Parametric and Non-parametric Tests Detect Significant Components in Power Spectrum. Notice that the non-normal distribution of the raw autospectra (top) leads to difference between the mean and median based estimates of auto-spectrum (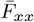 and 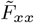). Both the parametric and non-parametric tests detect the significant bio-signal components (bottom) up to about 400-500Hz (the amplifier filter’s cut-off frequency) with the non-parametric test saturating due to finite sample size at lower frequencies with stronger effects, while performing better at detecting weaker high-frequency components compared to to its parametric counterpart. Corrected *α* corresponds to Bonferroni correction with the number of frequencies. See section 4.1 for details of the tests and section 6.2 for details of the data used.

Figure 11 compared the spectral EEG power between the 2 electrodes *C*_3_ and *C*_4_ above the contralateral and ipsilateral motor cortices of the active ongoing motor task. Both the parametric and non-parametric tests based on the raw distribution of auto-spectra prove useful in detecting low-alpha, low and high beta and low gamma changes of spectral power. The minor difference between the non-parametric and parametric tests (including stronger detections by the non-parametric test) can be due to minor difference in stationarity and the components from multiple physiological processes.

**Figure 11.**
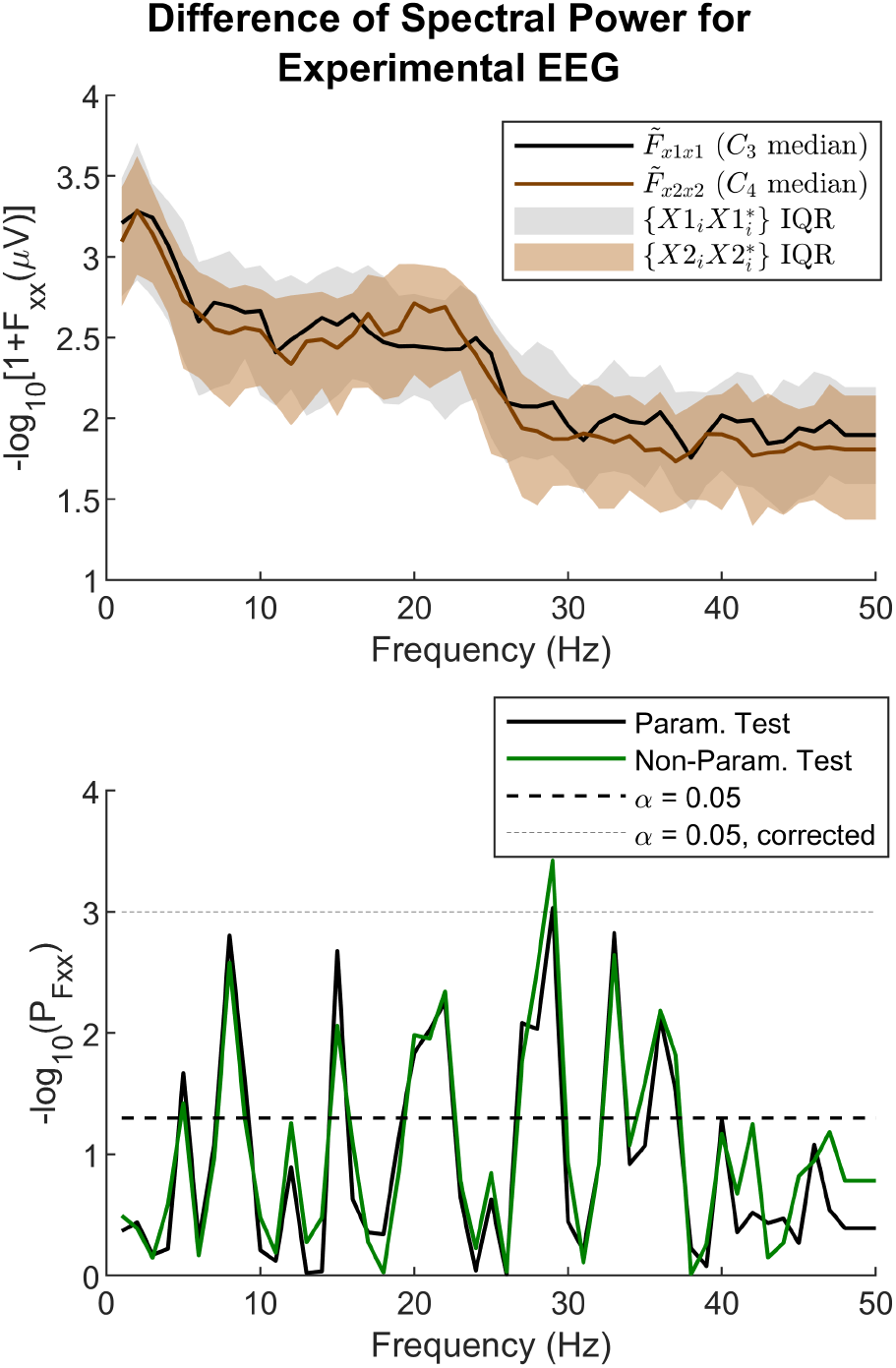
Experimental Data: Both Parametric and Non-parametric Tests Detect Significant Differences Between 2 Power Spectra. The 2 power spectra (top) correspond to EEG channels *C*_3_ and *C*_4_ over the left (contra-lateral) and right (ipsilateral) primary motor area of hand during an isometric pincher grip task. The p-values of both the parametric and non-parametric tests (bottom) detect the significant differences between the oscillation intensity in the 2 hemispheres. Corrected *α* corresponds to Bonferroni correction with the number of frequencies. See section 4.2 for details of the tests and section 6.2 for details of the data used.

#### 6.2.2 Coherence

Figure 12 shows the p-values corresponding to the traditional 1D statistical analysis of coherence using Equation (8), as well as the new 2D non-parametric test (Spatial Signed Rank). Both tests provide very comparable results. The non-parametric test provides stronger detection of synchronies in the beta and gamma band. The non-parametric detection of alpha band coherence is weaker. This can be due to the large transient alpha oscillations that are not dominant in the full distribution. The absolute value or the magnitude of coherence are both comparable with mean and median estimates. However, as the relationship between the coherence and p value is mediated by the number of epochs, there is always a bias term. This relationship is known for the parametric case, Equation (8), but is not available for the non-parametric case. The p-value spectrum therefore provides the same function and resolves these issues. The phase values corresponding to significant coherences show a range of non-trivial values using both the mean and median-based estimation, including a specific positive slope in alpha band and zero slop at high beta band.

**Figure 12.**
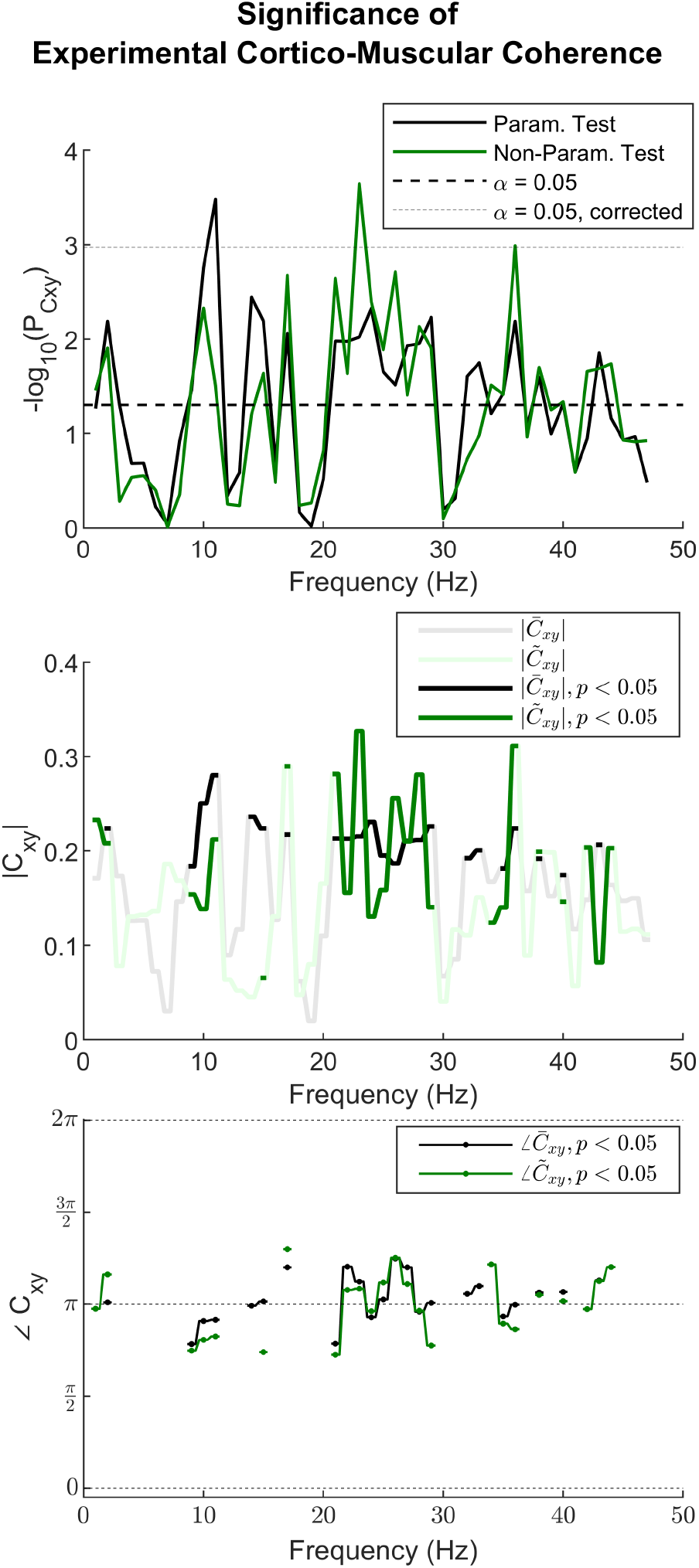
Experimental Data: Both Parametric and Non-parametric Tests Detect Significant Presence of Cortico-Muscular Coherence Between an EEG (*C*_3_) and an EMG (FDI) signals. **Top:** The p-values from the 1D parametric test on 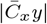 in Equation (8) and the 2D non-parametric Spatial Signed Rank test (section 5.1) show comparable detection of significant coherence. Corrected *α* corresponds to Bonferroni correction with the number of frequencies. **Middle:** The magnitude of coherence estimated using the mean 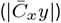 and median 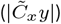, follow the p-values. **Bottom:** The phase of coherence estimated using the mean 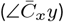 and median 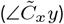 show close correspondence. See section 5.1 for details of the tests and section 6.2 for details of the data used.

Figure 13 show 2 different corticomusular EEG-EMG coherence measures: between *C*_3_-FDI and between *C*_4_-FDI. The median based 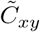 for both pairs show differences betwen these 2 channel pairs. The existing statistical techniques (parametric) do not provide direct comparisons of these measures at the level of individual subjects. The statistical trick in equation (20) provides a solution for an approximate inference between 2 pairs based on their individual p-values. This 1D comparison of magnitude (chosen to be based on *p*_1_ and *p*_2_ from non-parametric 1-sample tests) detects the changes in coherence strengths which is especially strong for low coherence values. The 2D non-parametric test based on spatial ranks detects changes in amplitude and especially the phase. A significant difference in the phase of synchrony in meaningful only if each of the signal pairs show a significant coherence in 1-sample tests, and, additionally, the 2-sample test yields significant differences. The certainty about the phase difference can be achieved if the 1D test of magnitude is non-significant, or alternatively by using a circular Mann-Whitney U-test (section 5.2.1) can confirm this.

**Figure 13.**
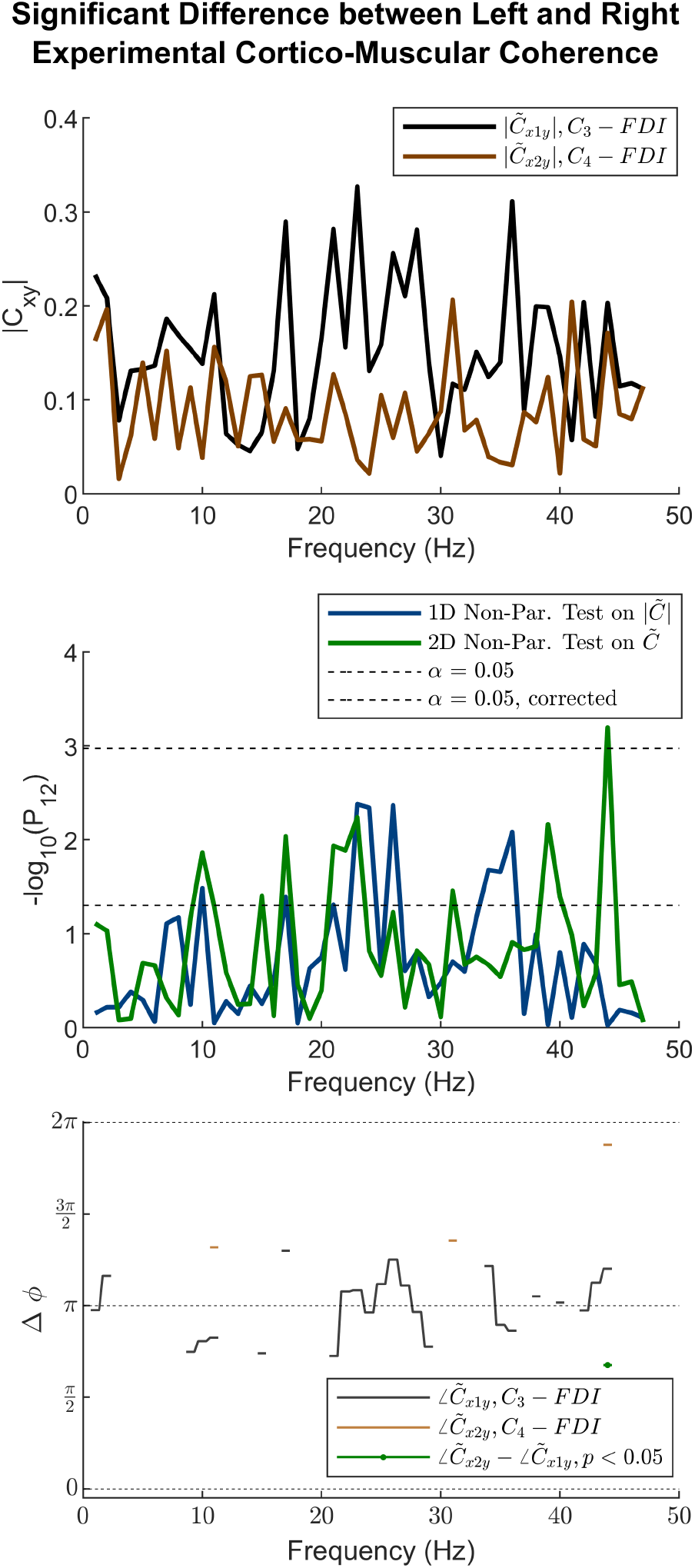
Experimental Data: Both Parametric and Non-parametric Tests Detect Significant Difference between Cortico-Muscular Coherence of 2 Signal Pairs. **Top:** The magnitude of coherence estimated using the median 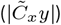, show differences between the cortico-muscular coherence in the contralateral and ipsilateral EEG electrodes and FDI muscle. **Middle:** The p-values from the 1D parametric test on 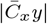 using the statistical trick for comparing 2 1-sample p-values (section 5.2.2) and the 2D non-parametric Spatial Rank test (section 5.2) both show frequency bands (especially the *β* band) with maximum differences. Corrected *α* corresponds to Bonferroni correction with the number of frequencies. **Bottom:** The phase of coherence estimated using the median 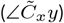 when the coherence is significant, is shown for *x*_1_(*C*_3_) and *y*(*FDI*), as well as for *x*_1_(*C*_4_) and *y*(*FDI*). The phase difference between 2 signal pairs is only plotted when both the individual pairs show significant difference, and when the 2D 2-sample non-parametric test shows significant difference. See section 5.2 for details of the tests and section 6.2 for details of the data used.

## 7 Discussion

### 7.1 The New Perspective and Methods

The new approach for spectral analysis takes into account the statistical distribution of the raw auto- and cross-spectra, enabling working with non-parametric estimates such as median and rank-based statistical tests. This approach which we previously introduced (Dukic et al., 2017) is considerably different from the established parametric methods on spectral analyses of time series.

The approach to conduct the test on the data in 2D complex plane is a novel direction that can be achieved reasonably only by non-parametric tests (given the complex distribution of the complex data). This approach takes into account both the magnitude and phase of coherence. This enables the 1-sample testing of the coherence magnitude non-parametrically which would not be possible using existing methods. Importantly, this 2D approach enables the 2-sample comparison of two coherence levels. This would not be possible otherwise, except only for magnitude using the statistical trick in section 5.2.2. Importantly these tests are applicable in individual subjects.

### 7.2 Pros of Cons of Non-Parametric Spectral Statistics

Similar to 1D non-parametric tests based on rank statistics such as Wilcoxon’s Signed Rank and Mann-Whitney’s U test, the non-parametric 2D tests based on spatial signed ranks or ranks afford robustness against artefacts and outliers. They are applicable on arbitrary distributions. Theoretically, and as demonstrated by simulations they may not be as effective or powerful as parametric counterparts. Nevertheless in practical situations as shown in the real data, they provided very acceptable performance.

The non-parametric methods help to avoid computationally expensive bootstrapping methods when parametric methods are not available or are not preferred, as they do not suffer from unwanted variability in estimations.

The magnitude of non-parametric coherence 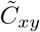 is directly related to the p-values from the tests. However, unlike the parametric estimates, a closed-form solution is still unknown or does not exist. This is not a concern as the bias of the coherence measure (a function of the number of epochs *L*) makes any interpretations dependent on the p-values associated to the coherence values. Non-parametric methods afford this p-value which can be plotted as −*log*_10_(*p*) spectrum and used as a more informative spectrum.

### 7.3 Applications

These developments and methods, while useful for broad range of neurophysiological research as well as in neuro-electromagnetic source imaging, are especially necessary for comparison of pathological conditions against healthy individuals. Importantly, the new methods allow analyses at individual subject level which would only be practical at the level of subject/patient levels.

### 7.4 Extension to Other Measures and Analyses

Given the general nature of the methods, they can be similarly used in an extended range of methods and analyses.

#### 7.4.1 Frequency and Time-Frequency Measures

The methods can be similarly applied on most of the complex-values frequency and time-frequency method. Examples include the spectra or coherence measures calculated using the Welch’s overlapping window (Terry & Griffin, 2008), spectral smoothing approach, or Multi-Taper techniques (Babadi & Brown, 2014; D. M. Halliday, Brittain, Stevenson, & Mason, 2018). Time-frequency representations based on complex values, including Short-Time Fourier Transform and Morlet wavelets (Boashash, 2003; Misiti, Misiti, Oppenheim, & Poggi, 2007) and their variations (Nasseroleslami et al., 2014; Mehrkanoon, Breakspear, Daffertshofer, & Boonstra, 2013) can be similarly tested using the non-parametric techniques.

#### 7.4.2 Partial and Multiple Spectra

Other derivatives of the spectral measures, such as the partial spectra, partial coherence (Lindsay & Rosenberg, 2011) and multiple coherence (D. M. Halliday & Rosenberg, 1999) can be similarly tested using non-parametric methods, as they are similarly expressed in the 2-D complex plane. Care should be exercised when the null hypotheses of these measures may vary in specific applications.

#### 7.4.3 Group Analysis

Combination of time series data from several subjects or several experimental sessions can be achieved using the pooled coherence (D.M. Halliday & Rosenberg, 1999) of all the raw data points in a group. Alternatively this can be achieved using the existing methods for combining the p-values such as Stouffer’s method (Stouffer, 1977; Westfall, 2014).

#### 7.4.4 High-Dimensional Statistics

Scientific inference from neural time series such as EEG and EMG signals may rely on analyses in high dimensions encompassing time, frequency and space/channels. The p-values from parametric and non-parametric analyses can be similarly corrected for multiple comparisons using the effective number of components (Nasseroleslami et al., 2014), random-field theory (Siegmund & Worsley, 1995; Mehrkanoon et al., 2014), or (adaptive) False Discovery Rate (Benjamini & Hochberg, 1995; Benjamini et al., 2006). Methods such as Empirical Bayesian Inference Efron, 2007; Nasseroleslami, 2018 applied to high-dimensional analyses of spectral measures (Nasseroleslami et al., 2017) affords the estimation of posterior probability and statistical power. Therefore, multiple comparison is straight-forward with new statistical methods.

#### 7.4.5 Source Analysis

The non-parametric statistical methods on spectral measures would greatly benefit the analysis of neural activity and connectivity in neuro-electromagnetic source analysis (Ramírez, Wipf, & Baillet, 2010). Given the application of spectral source anad connectivity analysis in healthy individuals and patient groups (Muthuraman et al., 2018; A. R. Anwar et al., 2016) and the crucial need to use robust unbiased measures of connectivity, the proposed methods can be of great utility.

#### 7.4.6 Future Directions

Future direction should work more on better ways to estimate the statistical power. This is automatically taken care of with some multi-variate methods such as empirical Bayesian inference. However, currently, the straight-forward method for finding power in low dimensional analysis is through non-null bootstrapping.

Paired 2-sample testing may be a possibility depending on the nature of analysis. This has not been adequately addressed to date. However, both parametric and non-parametric methods can be potentially used for such analyses and is expected to afford additional statistical power where applicable..

## 8 Conclusions and Recommendations

Non-parametric statistical analysis of spectral power and coherence can be used in several practical situations on 1-sample and 2-sample problems. In addition to the robustness afforded by non-parametric measures, based on rank or median, these statistical techniques are not restricted by parametric assumptions in traditional analysis and can be used in broader ranges of comparisons in neural signal analysis. In addition to rank-based 1-sample and 2-sample tests for comparing spectral power, the 2D Spatial Signed Rank and Spatial Rank Tests are the recommended tests for assessing the significance of coherence in 1-sample and 2-sample problems.

## Acknowledgement

The support of students and staff at Trinity College Dublin, the University of Dublin is highly appreciated.

This work was supported by Irish Research Council (Government of Ireland Postdoctoral Research Fellowship GOIPD/2015/213 to B.N.), Science Foundation Ireland (award SFI/16/ERCD/3854), and Health Research Board of Ireland (award HRA-POR-2013-246).

